# Relevance of Circulating Hybrid Cells as a Non-Invasive Biomarker for Myriad Solid Tumors

**DOI:** 10.1101/2021.03.11.434896

**Authors:** Matthew S. Dietz, Thomas L. Sutton, Brett S. Walker, Charles E. Gast, Luai Zarour, Sidharth K. Sengupta, John R. Swain, Jennifer Eng, Michael Parappilly, Kristen Limbach, Ariana Sattler, Erik Burlingame, Yuki Chin, Austin Gower, Jose L. Montoya Mira, Ajay Sapre, Yu-Jui Chiu, Daniel R. Clayburgh, SuEllen J. Pommier, Jeremy P. Cetnar, Jared M. Fischer, Jerry J. Jaboin, Seunggu J. Han, Kellie J. Nazemi, Rodney F. Pommier, Kevin G. Billingsley, Brett C. Sheppard, V. Liana Tsikitis, Alison H. Skalet, Skye C. Mayo, Charles D. Lopez, Joe W. Gray, Gordon B. Mills, Zahi Mitri, Young Hwan Chang, Koei Chin, Melissa H. Wong

## Abstract

**Abstract:** Metastatic progression defines the final stages of tumor evolution and underlies the majority of cancer-related deaths. The heterogeneity in disseminated tumor cell populations capable of seeding and growing in distant organ sites contributes to the development of treatment resistant disease. We recently reported the identification of a novel tumor-derived cell population, circulating hybrid cells (CHCs), harboring attributes from both macrophages and neoplastic cells, including functional characteristics important to metastatic spread. These disseminated hybrids outnumber conventionally defined circulating tumor cells (CTCs) in cancer patients. It is unknown if CHCs represent a generalized cancer mechanism for cell dissemination, or if this population is relevant to the metastatic cascade. Herein, we detect CHCs in the peripheral blood of patients with cancer in myriad disease sites encompassing epithelial and non-epithelial malignancies. Further, we demonstrate that in vivo-derived hybrid cells harbor tumor-initiating capacity in murine cancer models and that CHCs from human breast cancer patients express stem cell antigens, features consistent with the ability to seed and grow at metastatic sites. Finally, we reveal heterogeneity of CHC phenotypes reflect key tumor features, including oncogenic mutations and functional protein expression. Importantly, this novel population of disseminated neoplastic cells opens a new area in cancer biology and renewed opportunity for battling metastatic disease.

**Simple Summary:** There is an incomplete understanding of circulating neoplastic cell populations and the fundamental mechanisms that drive dissemination, immune evasion, and growth —all critical information to more effectively prevent and treat cancer progression. A novel disseminated tumor cell population, circulating hybrid cells, are detected across many cancer types and carry functional tumor-initiating properties. Additionally, circulating hybrid cells are found at significantly higher levels than conventionally defined circulating tumor cells. Our study demonstrates that neoplastic hybrid cells harbor phenotypic and genetic characteristics of tumor and immune cells, display stem features, and are a generalizable phenomenon in solid tumors. Circulating hybrid cells therefore have relevance as a novel biomarker and open a new field of study in malignancy.

## 1. Introduction

Metastatic disease contributes to over 90% of cancer-related deaths [1]. The disparity in survival between early- and late-stage disease reflects a knowledge gap in the biology underlying metastasis, such as the identity of tumor cells prone to disseminate and functional insight into their seeding of metastatic tumors. To successfully navigate the metastatic cascade, tumorigenic cells must invade surrounding tissue, intravasate into circulation, seed distant sites, and establish a permissive microenvironment for colonization and growth [2–4]. Among these events, the mechanisms that allow tumor cells to escape into circulation and survive are arguably the least understood. Direct investigation of tumor cells in circulation is hampered by their rarity, however it remains clear they provide an opportunity to better understand metastatic progression and potentially impact patient outcomes.

Detection of conventionally-defined circulating tumor cells (CTCs) in metastatic cancer patients provided the first direct evidence for tumor cell dissemination [5]. Distinct from other cells in circulation, CTCs express tumor proteins [e.g. cytokeratin (CK), epithelial cell adhesion molecule (EpCAM), and E-cadherin (ECAD) in epithelial malignancies] but lack leukocyte epitopes (e.g. CD45). CTCs are reported in many disease sites, including breast, colorectal, lung, pancreatic, and prostate cancer [*6–10*], as well as non-epithelial cancers, such as glioblastoma [11] and melanoma [12]. The tumor-based origin of CTCs is supported by their mutual expression of nucleic acids, protein and cellular features. However, initial excitement for the utility of CTCs has been tempered by their rarity, highlighted by just 5 CTCs/7.5 mLs of blood in the clinical presence of high tumor burden [7]. Moreover, CTC enumeration has failed to inform therapeutic decision making, especially in early-stage disease [13].

Our group reported a novel tumor-derived cell population detected in peripheral blood of patients with pancreatic ductal adenocarcinoma (PDAC) [14]. Circulating hybrid cells (CHCs) express neoplastic and immune cell functional attributes and are readily identifiable by their co-expression of tumor and leukocyte proteins. Our previous studies identified cell fusion as a major pathway for hybrid generation [14–17], though other described mechanisms can also result in tumor cells with dual expression states, including immune mimicry [18], developmental gene expression [19] and exosome-mediated protein/mRNA expression [20]. Focusing on cell fusion, our studies have rigorously demonstrated that neoplastic-immune cell hybrids retain functional genotypic and phenotypic properties from both parent cells based on culture assays, murine models of tumorigenesis, and human patient samples [14–16]. *In vitro*- and *in vivo*-derived fusion hybrids displayed enhanced motility, invasiveness and growth at metastatic sites, indicating they play a central role in the progression of metastatic disease [14]. In patients with pancreatic cancer, CHCs significantly outnumbered CTCs at every stage of disease, with levels capable of providing a prognostic indicator for overall survival [14]. Further, detection of CHCs with ECAD expression correlates with PDAC nodal staging [21], indicating subpopulations of CHCs with differential protein expression may provide a predictive biomarker. However, as a newly defined population, the functional tumor-initiating capacity and expression of stem cell attributes of CHCs is unexplored.

Tumors contain significant cellular heterogeneity and select populations exhibit key attributes needed to successfully spread to distant sites. Presumably, generation of neoplastic hybrid cells supports the evolution of cells with capabilities to both migrate and initiate tumors. The concept that neoplastic cells with stem cell properties represent a source for metastatic spread and treatment resistance is strongly supported by the literature [22–24]. Not only do neoplastic cells expressing stem cell surface antigens harbor tumorigenic potential, their levels within a tumor negatively correlate with patient survival in multiple cancer types [25–27]. In breast cancer, tumor-initiating cancer stem cells express CD44, a transmembrane receptor for hyaluronic acid that is a known effector of metastasis, and maintain low levels of CD24, a sialoglycoprotein that facilitates a number of important signaling networks in development [28–32]. This functional cell surface identity extends to other solid organ malignancies including PDAC, prostate cancer, colon cancer, hepatocellular carcinoma, melanoma, and glioma [33, 34]. Though rare, CTCs with tumorigenic ability mirror the expression of cancer stem cells, CD44^+^/CD24^lo^ [35, 36], and supports their prognostic value [37–40]. Based upon the metastatic potential of disseminated tumor cells, we sought to specifically investigate the prevalence of CHCs across numerous solid organ malignancies, identify sub-populations with stem cell markers, and evaluate conserved phenotypes between CHCs and tumor tissue.

Herein, we establish that CHCs represent the principal tumor-derived cell in circulation across a wide spectrum of solid organ malignancies, including epithelial and non-epithelial cancers. Using murine models and human patient samples, we provide evidence that hybrid cells possess tumor-initiating behaviors and characteristics that indicate their potential importance in metastatic disease. Furthermore, we reveal CHCs share phenotypic similarities with tumor tissue and reflect the cellular heterogeneity of neoplastic disease. With this evidence, we postulate that CHCs are effectors of metastatic spread and have translational value as a non-invasive analyte across cancer types.

## 2. Results

### CHCs are the dominant circulating tumor-derived cell in myriad cancers

Dissemination of neoplastic cells into circulation is a key step in metastatic progression. Therefore to determine if hybrid formation and escape is generalizable across cancer types, we evaluated peripheral blood for CHCs from patients with 14 different epithelial or non-epithelial malignancies, including ampullary adenocarcinoma, breast adenocarcinoma, ovarian carcinoma, cholangiocarcinoma, colon adenocarcinoma, esophageal cancer, high grade glioma (pediatric & adult), head and neck squamous cell carcinoma, pancreatic ductal adenocarcinoma, pancreatic neuroendocrine tumor, prostate adenocarcinoma, rectal adenocarcinoma and uveal melanoma (Table 1). A standard ficoll-density gradient facilitated the isolation of peripheral blood mononuclear cells (PBMCs) and downstream CHC detection. Circulating tumor-derived cells were identified by protein expression of canonical tumor markers [Epithelial: CK^+^; Uveal melanoma: NKI beteb^+^; Glioma: glial fibrillary acidic protein (GFAP^+^) [41]; Pancreatic neuroendocrine tumor (PNET): chromogranin A (CHGA^+^), synaptophysin (SYP^+^) [42]] using two different platforms, flow cytometry and fluorescence microscopy. Expression of the pan-leukocyte antigen CD45 provided the distinction between CTCs and CHCs, where CHCs were identified as cells co-positive for both tumor protein and CD45 (Figure 1, Figure S1). Flow cytometry facilitated robust quantitative analyses (Figure 1B, 1C), while microscopy provided visual confirmation of protein expression and their relevant cellular localization (Figure 1D, 1E, Figure S2). Leveraging the collective power of these two platforms, we identified CHCs in every cancer we analyzed, including glioma, which is reported to seldomly disseminate outside of the central nervous system [43]. In addition, we identified a significantly higher number of CHCs in each disease site relative to healthy subjects (*p* < 0.00001 for all, except PDAC, ECA, Adult Glio *p* < 0.0001, and HNSCC *p* < 0.05), and found that CHCs outnumbered CTCs in all cancer types (Figure 1, range of *p* < 0.001-0.0001). Despite differences in age, sex, tumor burden, and prior treatment status, across cancers, the heterogeneous patient population (Table S1) harbored higher levels of CHCs than CTCs.

**Table 1.** Patient Clinicopathologic Characteristics by Disease Site

| Disease Site | Number of Patients | Age Range | Disease State |  |  | Treatment Naïve (%) |
| --- | --- | --- | --- | --- | --- | --- |
|  |  |  | Local | Regional | Metastatic |  |
| Ampullary Adenocarcinoma | 5 | 56-77 | 2 | 3 | 0 | 5 (100) |
| Breast Adenocarcinoma | 5 | 51-73 | 1 | 2 | 2 | 4 (80) |
| Cholangiocarcinoma | 5 | 42-81 | 2 | 1 | 2 | 5 (100) |
| Colon Adenocarcinoma | 5 | 33-65 | 0 | 0 | 5 | 1 (20) |
| Esophageal Cancer | 5 | 52-75 | 3 | 2 | 0 | 0 (0) |
| Adult High-Grade Glioma | 5 | 31-74 | 5 | 0 | 0 | 2 (40) |
| Pediatric High-Grade Glioma | 5 | 4-15 | 5 | 0 | 0 | 1 (20) |
| Head & Neck SCC | 5 | 57-70 | 0 | 5 | 0 | 5 (100) |
| Non-Small Cell Lung Cancer | 5 | 41-74 | 0 | 0 | 5 | 2 (50) |
| Ovarian Carcinoma | 5 | 60-76 | 0 | 4 | 1 | 1 (20) |
| Pancreatic Adenocarcinoma | 5 | 56-76 | 1 | 2 | 2 | 1 (20) |
| Pancreatic NET | 5 | 55-88 | 4 | 1 | 0 | 5 (100) |
| Prostate Adenocarcinoma | 5 | 57-72 | 5 | 0 | 0 | 5 (100) |
| Rectal Adenocarcinoma | 5 | 47-67 | 0 | 0 | 5 | 0 (0) |
| Uveal Melanoma | 5 | 56-71 | 4 | 0 | 1 | 5 (100) |
| Total | 75 | 4-88 | 32 (43%) | 20 (27%) | 23 (31%) | 42 (56%) |
| Abbreviations: CHC=Circulating Hybrid Cell; SCC=Squamous Cell Carcinoma; NET=Neuroendocrine Tumor |  |  |  |  |  |  |

**Figure 1.**
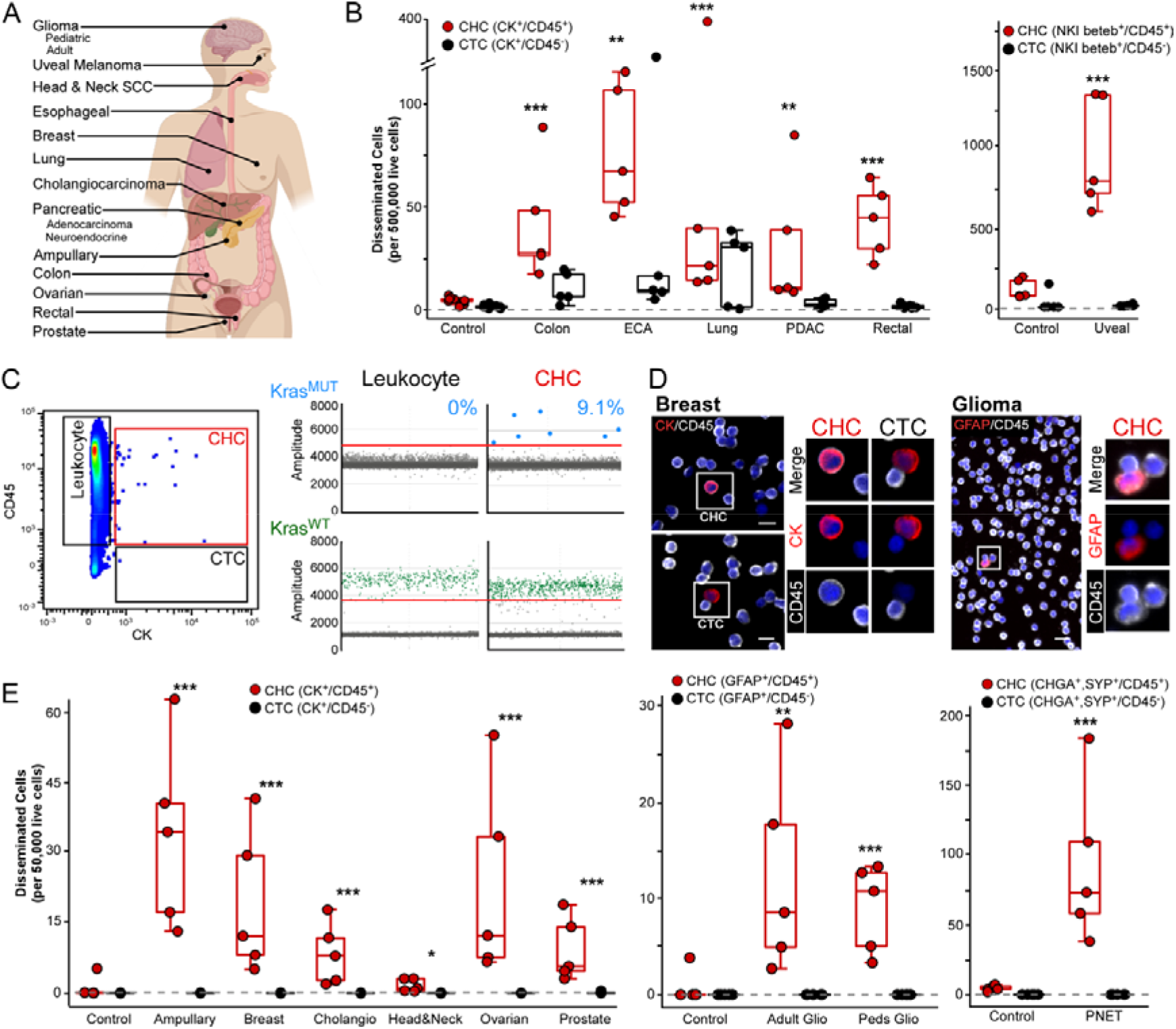
Circulating Hybrids Cells out number. Circulating Tumor Cells in myriad of human cancer disease sites. Circulating Hybrids Cells out number Circulating Tumor Cells in myriad of human cancer disease sites. A) Anatomical location of 14 cancers evaluated for disseminated CTCs and CHCs. B) Flow cytometry analysis of peripheral blood from patients with advanced stage colon, esophageal, lung, pancreatic and rectal adenocarcinomas as well as uveal melanoma (n = 5 patients per disease site) demonstrates CHCs (mean value, red bar) present at significantly higher levels compared to CTCs (mean value, grey bar) and healthy controls (mean value, n=10). C) Representative flow cytometry scatter plot of cytokeratin vs CD45 expression, CHCs (red box), CTCs (gray box) full gating schema in fig. S1. Isolation of CHCs from pancreatic adenocarcinoma identifies KRAS mutation in 9.1% of CHCs using ddPCR. D) In situ immunofluorescence microscopy analysis of peripheral blood cells analyzed for detection of glial fibrillary acid protein (GFAP; red) and CD45 (white) expression in pediatric high-grade glioma and cytokeratin (CK; red) and CD45 (white) in breast adenocarcinoma, identify CHCs and rare CTCs. Scale Bars 10 and 20 μm, respectively. E) Immunofluorescence microscopy analysis of peripheral blood from patients (n=5 per disease site) with ampullary carcinoma, breast, cholangiocarcinoma, head and neck squamous cell carcinoma, ovarian, prostate, high grade glioma (adult and pediatric) and pancreatic neuroendocrine tumors reveals CHCs (mean value, red bar) are present at significantly higher levels compared to CTCs (mean value, grey bar) and healthy controls. \*\*\**P*< 0.00001; \*\**P*< 0.0001; \**P*<0.05

To further establish that CHCs derive from tumor tissue, we analyzed CHCs for the presence of known oncogenic mutations. Oncogenic *KRAS* mutations are implicated in the malignant transformation of pancreatic epithelia and are almost ubiquitously present in PDAC tumors [44, 45]. To define the oncogenic identity of CHCs at the genomic level, we focused our analysis on PDAC-derived CHCs. Using fluorescence-activated cell sorting (FACS), we isolated normal leukocytes (7000; CD45^+^/EpCAM^−^/ECAD^−^/CD49c^−^) and CHCs (580; CD45+/[EpCAM/ECAD/CD49c]^+^). CTCs were too rare for capture by FACS. Using digital droplet polymerase chain reaction (ddPCR), we screened samples for wild type *KRAS* and seven common oncogenic *KRAS* mutations [45, 46]. We identified that up to 9% of the isolated CHCs harbored an oncogenic *KRAS* allele while the leukocyte fraction only expressed wild type *KRAS* (Figure 1C), not only indicating that CHCs derive from the primary tumor, but also retain key genomic drivers of cancer progression.

### Hybrid cells harbor tumor-initiating properties

Retention of oncogenic driver mutations in disseminated tumor cell populations supports their relevance in metastatic disease progression. However, evidence of the capacity for CHCs to seed and grow metastatic tumors has not been previously explored. Our prior studies demonstrated that fusion-hybrids derived from allografted B16F10 melanoma cells are detectable in both primary and metastatic sites, with greater tumorigenicity than unfused cancer cells [14]. However, as they are derived from immortalized cell lines uniformly transformed and selected for their proliferative properties, this model is suboptimal to assess tumor-initiating potential. In order to more accurately investigate the contribution of hybrid cells to metastatic progression, we generated a model for isolating fusion hybrids derived from autogenous malignancy. Here, we engaged the murine mammary tumor model (mouse mammary tumor virus-polyoma middle tumor-antigen; MMTV:PyMT), which develops mammary carcinoma [47]. We genetically marked tumor cells with red fluorescent protein (RFP) by crossing the MMTV-PyMT mouse onto a CAG-RFP background, yielding MMTV-PyMT-RFP mice. Harvested RFP^+^ mammary tumor cells were dissociated into single cells and injected into the mammary fat pad of Actin-green fluorescent protein (GFP) transgenic mice, generating tumors designated as MMTV-PyMT-RFP into-GFP (Figure 2A). In this model, co-expression of RFP and GFP identified tumor hybrid cells.

**Figure 2.**
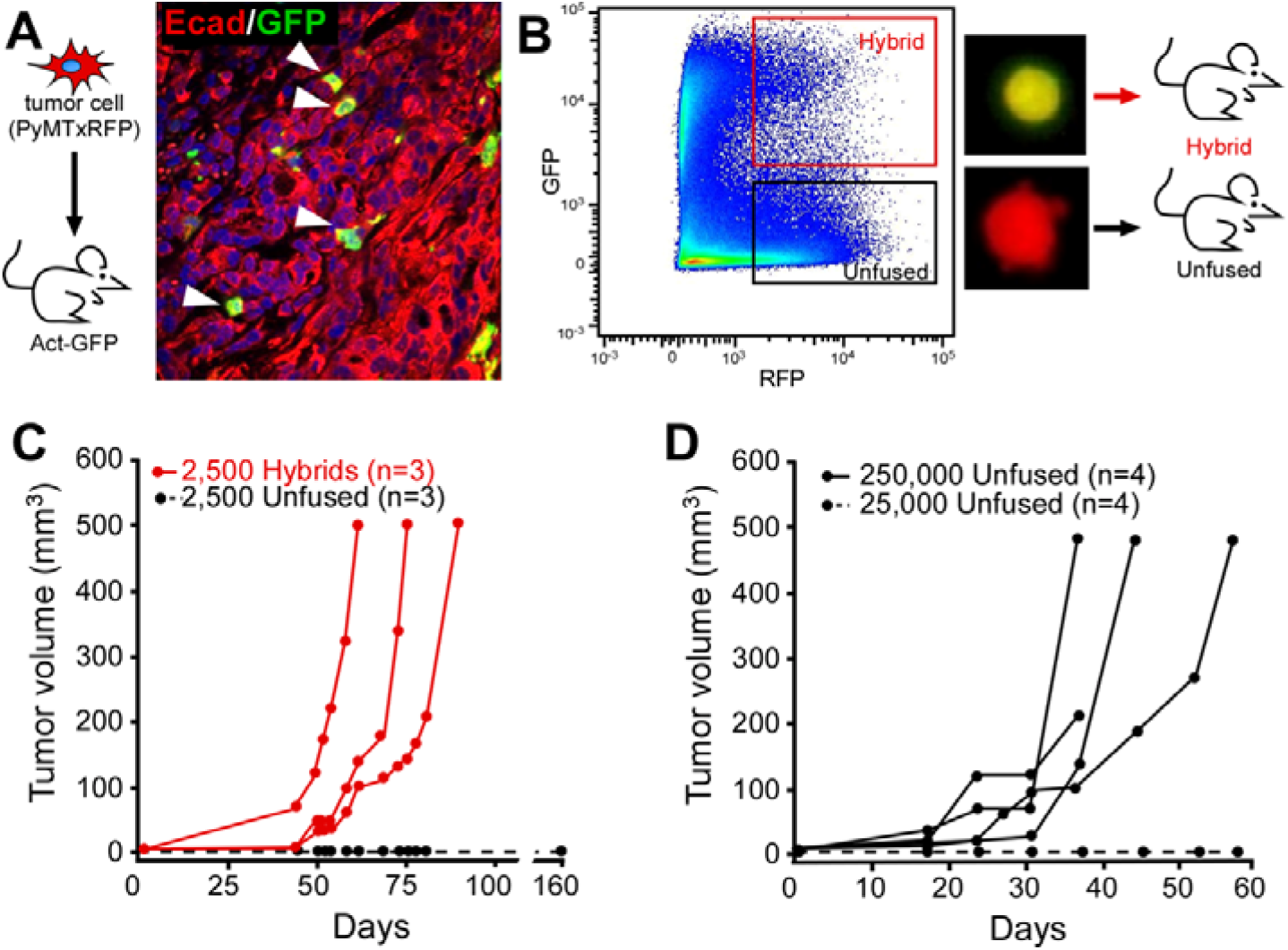
Murine hybrid cells harbor tumor-initiating capacity. Murine hybrid cells harbor tumor-initiating capacity. A) Mammary carcinoma cells from PyMT:RFP mice injected into GFP-expressing recipients. Resulting tumors express E-cadherin (red) and Actin:GFP (green) and were counterstained with Hoechst dye (blue). Arrowhead denotes co-positive hybrid cells. B) Representative FACs plot of tumor-derived hybrid cells (GFP+/RFP+) and unfused tumor cells (RFP+/GFP−) from dissociated tumor and injected into syngeneic wt recipient mice. C) Twenty-five hundred FACS-isolated cells were injected into recipient mice (hybrid cells, red line) and (unfused cells, dashed black line) and temporally monitored for growth. D) Injection of 250,000 unfused tumor cells developed tumors in recipient mice (black line), while 25,000 cells did not grow (dashed line).

To assess the tumor-forming capacity of hybrid cells, we FACS-isolated hybrid cells (RFP^+^/GFP^+^) and unfused tumor cells (RFP^+^/GFP^−^) from MMTV-PyMT-RFP into-GFP tumors (Figure S3). We then independently injected each cell type into the mammary fat pad of secondary wild type recipient mice (2500 cells per animal; Figure 2B). Neoplastic-derived hybrids supported rapid tumor growth, whereas unfused cancer cells did not generate tumors (Figure 2C). To further investigate relative tumor-initiating capacity, we used a limiting-dilution assay. We found that to generate tumors in all mice, two orders of magnitudes more unfused cancer cells were required compared to the number of hybrids (Figure 2D). The potent tumor-initiating capacity of hybrid cells is consistent with functional stem cell capacities. Cancer stem cell phenotypes, which are widely identified in human tumors, are associated with increased tumorigenic potential compared to their non-stem cell counter parts, have been linked to CTC phenotypes [32, 48–50]. The functional resemblance between fusion-hybrids and cancer stem cells necessitates investigation as whether human CHCs similarly harbor stem characteristics.

### Human breast cancer CHCs display stem cell phenotypes

To investigate stem cell attributes of human CHCs, we leveraged the well-described immunophenotype of tumorigenic breast cancer stem cells identified by CD44^+^/CD24^lo^ expression [28–32]. We evaluated peripheral blood specimens from a cohort of twenty-seven treatment-naïve patients with breast cancer representing all stages and subtypes (Table 2, Table S2). Using flow cytometry, PBMCs were interrogated for expression of EpCAM, CD45, CD31, CD44 and CD24 to identify CTC and CHCs with stem cell identity (Figure 3). Consistent with the findings from our pan-cancer evaluation (Figure 1), we identified greater numbers of CHCs than CTCs in all breast cancer patients (Figure 3A, Figure S4). Furthermore, CHCs display high proportions of stem cell characteristics. All twenty-seven patient samples harbored CD44^+^/CD24^lo^ CHCs while only 74.1% (n=20) had detectable CD44^+^/CD24^lo^ CTCs. When present, CD44^+^/CD24^lo^ CTCs were found at significantly lower level than CD44^+^/CD24^lo^ CHCs, with means of 1.0 and 45.3 per 500,000 live cells, respectively (*p* < 0.00001). Additionally, compared to the normal leukocyte populations (CD45^+^/EpCAM^−^), there was a higher proportion of CHCs with CD44^+^/CD24^lo^ expression (*p* < 0.01), indicating the high level of CHCs with stemness features does not simply reflect the normal distribution of leukocyte phenotypes. The expression of stem markers by CHCs further supports their relevance in metastatic progression. Coupled with their relative abundance compared to CTCs, these data suggest CHCs are a readily available liquid analyte to query features of solid-organ malignancies, particularly in early-stage disease. These results prompted our investigation into the extent to which CHCs reflect the discrete phenotypic features and diversity of neoplastic cells in cancer tissue.

**Table 2.** Clinicopathologic Characteristics of Breast Cancer Patients Analyzed for CHC Stem Cell Properties

| Cancer Subtype | Number of Patients | Age Range | Disease State |  |  | Treatment Naïve (%) |
| --- | --- | --- | --- | --- | --- | --- |
|  |  |  | Local | Regional | Metastatic |  |
| HR<sup>+</sup> / HER2<sup>-</sup> | 15 | 28-80 | 8 | 5 | 2 | 15 (100) |
| HR<sup>+</sup> / HER2<sup>+</sup> | 7 | 38-77 | 3 | 3 | 1 | 7 (100) |
| HR<sup>-</sup> / HER2<sup>-</sup> | 3 | 26-58 | 1 | 2 | 0 | 3 (100) |
| HR<sup>-</sup> / HER2<sup>+</sup> | 2 | 44-50 | 0 | 1 | 1 | 2 (100) |
| Total | 27 | 26-80 | 11 (41%) | 12 (44%) | 4 (15%) | 27 (100%) |

Abbreviations: CHC=Circulating Hybrid Cell; HR=Hormone Receptor; HER2=Human Epidermal Growth Factor Receptor 2

**Figure 3.**
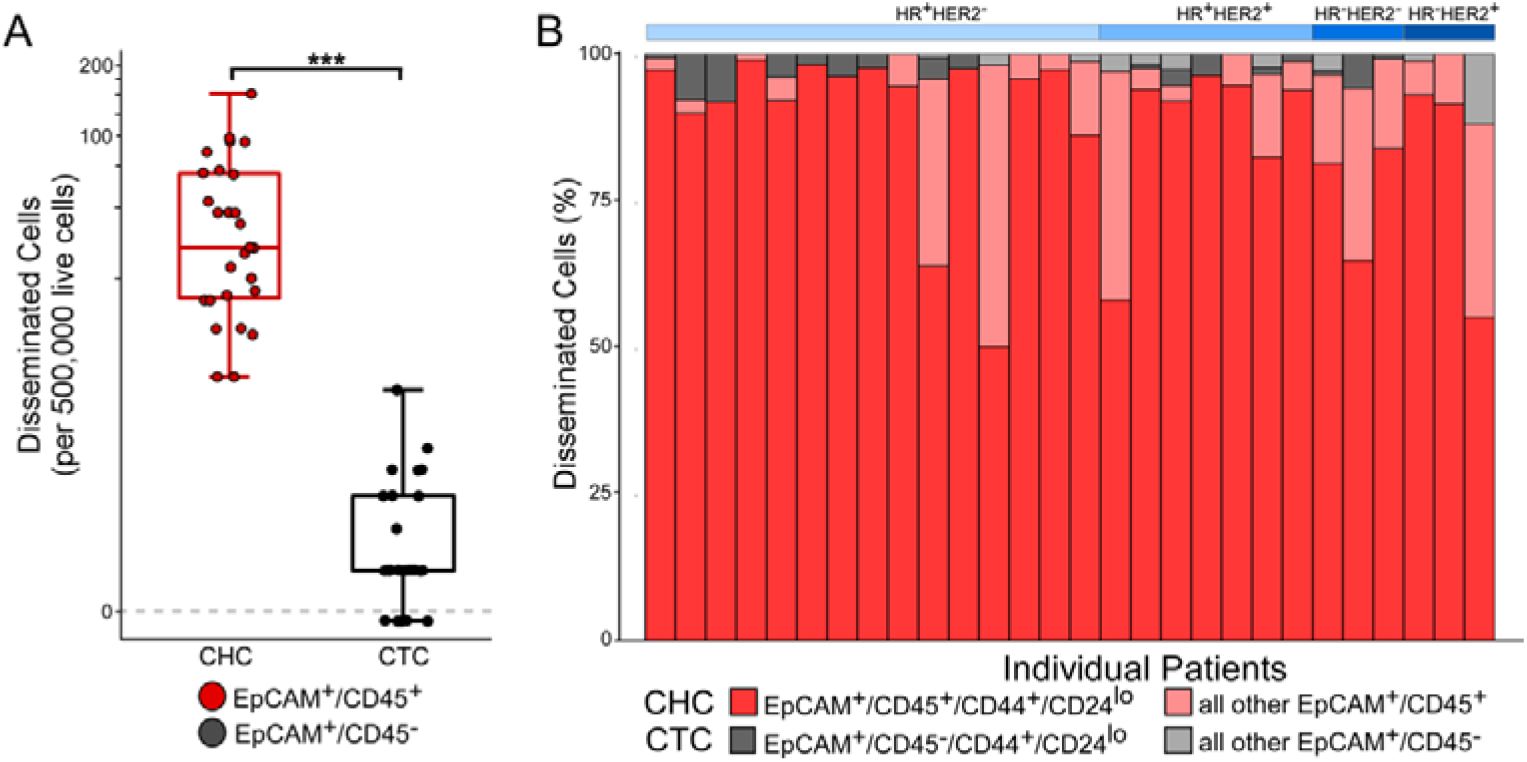
CTC and CHC stem cell antigen expression in breast cancer patients. CTC and CHC stem cell antigen expression in breast cancer patients. A) Flow cytometric analysis of peripheral blood from 27 untreated patients across breast cancer subtypes. Box whisker plot of CHCs (EpCAM+/CD45+; red) and CTCs (EpCAM+/CD45-; gray). CHCs levels are higher than CTCs at significant levels (p<0.0001). B) Depiction of percentages of CHC (red scale) and CTCs (gray scale) with stem cell identify (i.e. CD44+/CD24lo).

### Tumors disseminate a heterogeneous population of CHCs

The relative prevalence of CHCs and their suitability for liquid biopsy [51] provides an exciting and unexplored opportunity to query their diagnostic, prognostic, or predictive utility for longitudinal care of cancer patients. One major challenge in the management of solid tumors is the vast heterogeneity that promotes variable or incomplete response to available therapeutics. Standard tissue biopsies may fail to capture the full spectrum of tumor heterogeneity or miss cells that would otherwise inform known treatment vulnerabilities and resistances. Further, repeat tumor sampling is fraught with logistical challenges and risks patient harm. CHCs are detectable in sufficient numbers to facilitate more complete tumor analyses (Figure 1C), which could offer insight into tumor phenotyping and evolution over the course of a patient’s treatment. To explore how CHCs reflect the phenotypic features and diversity within the tumor, we analyzed tumor biopsy specimens and corresponding CHCs from a patient with refractory metastatic breast cancer while receiving palliative chemotherapy (Figure 4). The patient underwent an initial biopsy (BX1) at time of study enrollment and started monotherapy with the poly (ADP-ribose) polymerase (PARP) inhibitor, olaparib. A second biopsy (BX2) was obtained a month later. This evaluation aimed to provide feasibility data to determine if real time phenotypic changes within tumor cells from the biopsy or in CHCs reflected response to treatment.

**Figure 4.**
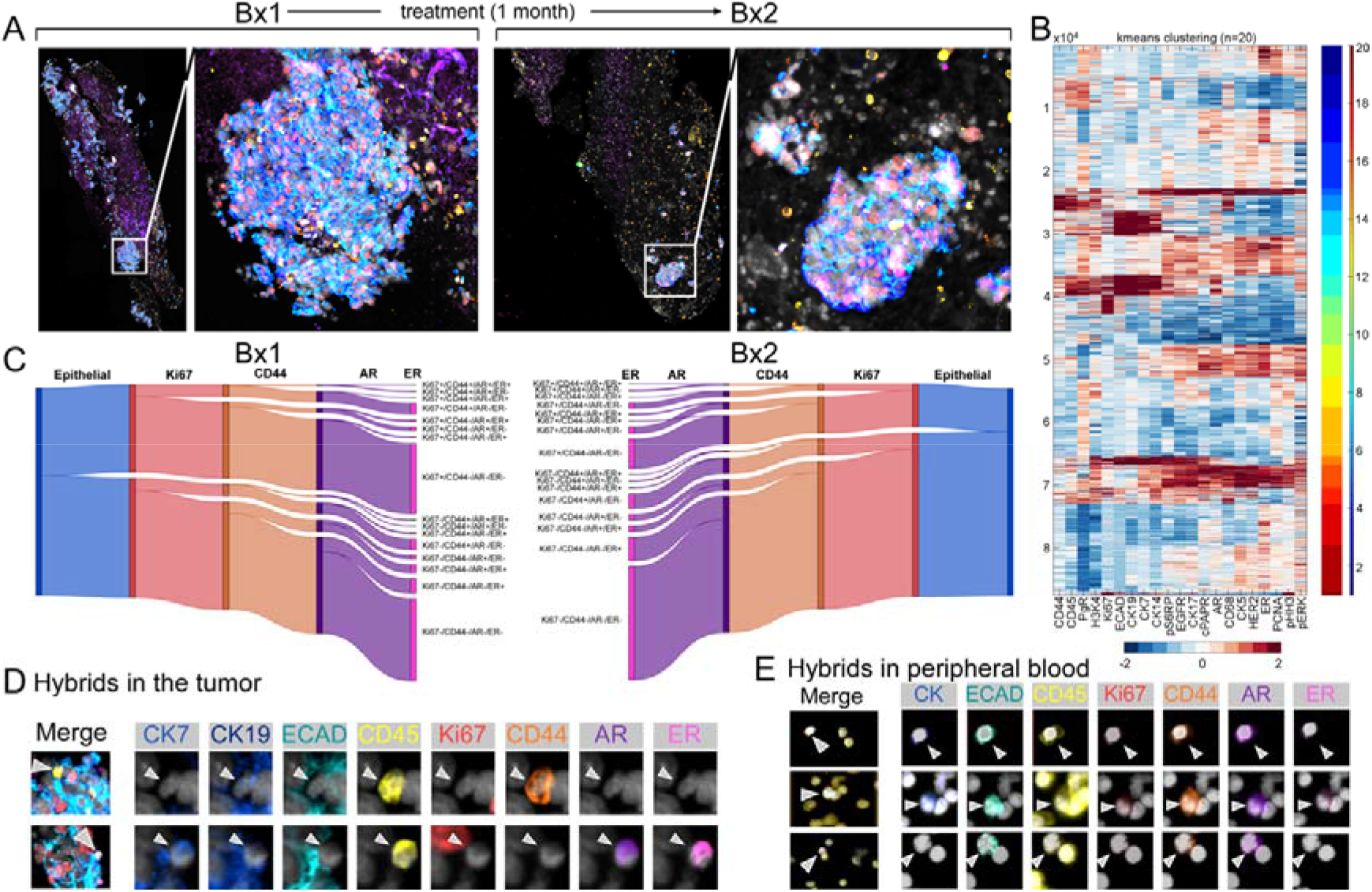
Multiplexed phenotypic analysis of tissue biopsy and CHCs from a subject with relapsed refractory breast cancer. Multiplexed phenotypic analysis of tissue biopsy and CHCs from a subject with relapsed refractory breast cancer. A) Cyclic Immunofluorescence microscopy evaluation of tumor biopsy tissue before and during treatment reveals hot spots of viable disease. B) Unsupervised clustering (k-means clustering with the number of clusters (n=20)) of cellular immunophenotype identified distinct clusters and reveal tumor heterogeneity. C) Sankey diagram of tumor phenotypes [proliferative (Ki67), stem (CD44), hormonal status (AR, HR)] quantifies heterogeneity of epithelial marker positive tumor cells. D) Representative images of heterogenous populations of hybrid cells expressing tumor epithelial (CK and/or ECAD) and immune (CD45) markers within biopsied tumor and E) peripheral blood mononuclear cells.

To identify treatment resistant disease, we spatially defined the tumor microenvironment and interrogated disease heterogeneity with a multiplexed cyclic immunofluorescence (cyCIF) on longitudinal hilar lymph node biopsies from a patient with metastatic triple-negative breast cancer (TNBC). CyCIF utilizes iterative staining and imaging cycles to facilitate the spatial resolution of >40 epitope-specific antibodies (Table S3) at the cellular level [52] facilitating resolution of various tumor features e.g. stromal, immune, epithelial and vascular compartments. We performed cyCIF on formalin-fixed paraffin-embedded (FFPE) tumor biopsies to phenotypically identify viable treatment-resistant disease hot spots, and gain insight into cellular heterogeneity and the complexity of the tumor’s ecosystem (Figure 4A). Extraction of fluorescent intensity patterns from multiplexed, segmented images allowed for single-cell unsupervised learning to probe tumor heterogeneity [53]. Using cellular features from BX1 and BX2, K-means clustering revealed discrete tumor cell populations based on clinically relevant protein expression (Figure 4B). To interrogate intratumoral and inter-biopsy heterogeneity we focused on the neoplastic epithelial compartment (ECAD^+^ and/or CK^+^ cells) to quantitatively describe the phenotypes of viable, treatment-resistant disease (Figure 4C). Within BX1 (pre-treatment), 42.5% of the epithelial compartment was Ki67^+^, with 14.9% of these cells also expressing CD44, suggestive of a proliferative stem cell phenotype. The distribution of hormone receptors, estrogen receptor (ER) and androgen receptor (AR), varied within this population, with 1.5% ER^+^/AR^+^, 1.4% ER^+^/AR^−^, and 6.5% ER^−^/AR^+^. After initiating therapy, only 20.6% of the malignant epithelia in BX2 were Ki67^+^ while the proportion of CD44^+^ cells remained similar at 14%, which may indicate a global tumor response to PARP inhibition but less effect on the proliferative stem population. The ER^+^/AR^+^ and ER^+^/AR^−^ cells remained consistent (1.2% and 0.6%, respectively), however we observed an increase in ER^−^/AR^+^ cells to 20% between BX1 and BX2. Importantly, AR-expression is associated with cellular proliferation and metastatic spread in TNBC [54], therefore the observed threefold increase in concentration of Ki67^+^/ER^−^/AR^+^ neoplastic cells under PARP inhibition likely indicates enrichment of a treatment-resistant proliferative cell population. In addition, we qualitatively identified hybrid cells within the tumor biopsy by the co-expression of CD45 and epithelial tumor proteins, ECAD and cytokeratins, including those with the treatment-resistant tumor phenotype (Figure 4D). Though only a qualitative analysis, it provides insight to the formative expression states of hybrid cells and further supports their relevance as a biologic mechanism contributing to tumor heterogeneity and disease evolution. With the identification of proliferative, treatment resistant disease within the tumor of this patient, we sought to investigate disseminated tumor cell populations from matched peripheral blood samples.

To investigate heterogeneity among tumor cells in circulation, we utilized cyCIF to interrogate PBMCs sampled at the time of BX2. To maximize phenotypic profiling and minimize cell loss, we performed two iterative staining cycles using antibodies to CD45, ECAD, panCK, AR, ER, Ki67 and CD44. Our analysis revealed a heterogeneous population of CHCs, including those with CD44 and AR expression (Figure 4E), which aligns with the phenotypes detected in the proliferating tumor compartment and tumor hybrids cells (Figure 4D). These data suggest that tumors disseminate a heterogeneous population of CHCs that reflect the overall phenotype of viable tumor, supporting the value of phenotypically profiling CHCs to anticipate disease evolution and treatment resistance.

## 3. Discussion

A cancer cell’s successful navigation of the metastatic cascade drives cancer lethality, highlighting the importance of understanding the functional biology of disseminated tumor cells. Recently, we identified a novel disseminated tumor cell population harboring both neoplastic and immune cell identities, and now establish their conserved functional phenotypes. Here, we demonstrated the presence of CHCs in patients across many epithelial and non-epithelial cancers, indicating that the generation of hybrids and their escape into peripheral blood is a ubiquitous and generalizable phenomenon in solid organ malignancies which warrant in-depth investigation. Although CHCs consistently outnumber CTCs across malignancies, the diversity and limited number of patients in each disease group prevents more robust exploration of the significance of their overall levels. However, as our prior analysis revealed CHC burden correlates with PDAC stage and patient survival, studies with larger cohorts should be pursued across cancer types to determine how CHC quantification might translate to clinical practice. The data presented here also indicates that phenotypic analysis of CHCs provides insight into tumor heterogeneity, and therefore may also hold prognostic value when queried for specific markers of interest (i.e. stem features or therapeutic targets). Further, as highlighted by the identification of PDAC-derived CHCs harboring *KRAS* mutations, CHCs hold considerable promise for use as a liquid biopsy to facilitate tumor profiling for clinically actionable oncogenic mutations. It is worth noting that the frequency of oncogenic *KRAS* mutations detected by ddPCR in CHCs from a single pancreatic cancer patient is similar to what has been reported for CTCs using the same method [55]. Although 93% of PDAC tumors harbor mutated *KRAS* genes [56], there is increasing evidence of the heterogeneity and distribution of *KRAS* mutations within individual tumors [56, 57]. The burden of *KRAS* mutations likely influences PDAC biology [58, 59], with worse survival reported in patients with ratios of mutant to wild type alleles >10% [57]. Selection for cells with more aggressive phenotypes, predilection for dissemination, or fusogenic potential may partially account for discrepancies seen between primary tumors and tumor-derived cells in circulation, and provide a rational basis for further investigation.

Disseminated neoplastic cells are effectors of tumor progression, yet the extent to which they reflect the spectrum of diversity among primary tumor cells remains unresolved. While it is possible that tumors randomly shed neoplastic cells into circulation, an active process may exist to mediate phenotype acquisition for escape from permissive tumor regions. To begin to unravel this question, we focused on disseminated cells in breast cancer patients and queried the functional state of CHCs. Using a tractable murine system to collect CHCs, we investigated the tumor-initiating capacity of hybrid cells and translated our findings to human disease by evaluating human CHCs for stem phenotypes. In a murine mammary tumor model, *autogenous* hybrid cells demonstrate the defining feature of stemness, namely growth *in vivo*, and with greater potency than unfused cancer cells. These findings suggest that hybrid identity independently supports their *in vivo* replicative potential and is reminiscent of stem cells, though the underlying mechanisms remains to be investigated. Our observation that classical breast cancer stem surface antigens (CD44^+^/CD24^lo^) [32–34] are enriched on breast cancer CHCs, suggests their tumor initiating potential. Additionally, it is interesting to note that CD44 is a known effector of homotypic macrophage fusion in experimental models [60], and its high expression in CHCs may indicate a previously unappreciated link between cancer stem cells and macrophage-cancer cell fusion. Beyond the ethos of CHC biology, focusing on the heterogeneity of disseminated tumor cells with stem phenotypes may expose cellular attributes necessary to navigate the metastatic cascade, such as homing to a premetastatic niche and the contributory features of proliferation, quiescence, senescence and cell cycle status. Moreover, interrogation of CHC and CTC phenotypes may illuminate evolving tumor heterogeneity to facilitate recognition of treatment resistant disease, and therefore be clinically leveraged as a liquid biopsy.

Cellular heterogeneity and the tumor microenvironment are at the forefront of translational research, as treatment failures are increasingly thought to evolve from resistant populations derived from neoplastic cell diversity. However, the extent to which circulating tumor-derived cells recapitulate the heterogeneity within tumor tissue remains unclear. We demonstrate that tumor tissue is spatio-temporally variable with a heterogeneous collection of tumor cells as reflected in the phenotypes of CHCs. Our comparative evaluation of tumor tissue and CHC phenotypes was performed with a metastatic hilar lymph node, as prior treatment precluded our ability to obtain primary breast tumor tissue. While the lack of primary tumor tissue is a limitation to our study, our findings highlight that tumor heterogeneity extends to metastatic sites and traditional biopsy methods are unlikely to illustrate the complete tableau of a patient’s disease. Additionally, our study showcases that neoplastic-immune cell hybrids are not limited to the primary tumor; however, we are cautious in generalizing the presence of hybrid cells within metastatic tumor tissue because of our limited sampling, the rarity of tumor hybrids and the qualitative nature of the analysis. Importantly, these findings do support the potential power of CHCs to aid comprehensive tumor analysis and monitor changes in disease. Indeed, our highly multiplexed immunofluorescence analyses of tumor and peripheral blood samples provide an unprecedented spatial resolution of cellular heterogeneity that allowed for identification of diverse populations of hybrid cells in tumors and in circulation. These observations, taken with the ubiquitous nature of CHCs in cancer and the robust tumor-initiating capacity of hybrid cells *in vivo*, suggest that CHCs may be a functioning population of cells relevant to the metastatic spread of cancer.

Only a minority of primary tumor cells functionally contribute to metastatic seeding. It is therefore unsurprising that phlebotomy may yield a more selective biopsy by enriching for biologically active tumor-derived cells. These advantages over standard tissue biopsy may provide an opportunity to monitor tumor evolution through serial blood draws. The abundance of CHCs relative to CTCs and the rarity of co-positive cells in healthy subjects supports their relevance in diagnostic and disease monitoring strategies. Further, the presence of oncogenic *KRAS* mutations in PDAC derived CHCs suggests their potential utility as a noninvasive analyte of disease biology. Finally, our data support the concept of the enriched liquid biopsy, as evidenced by our findings of robust stem populations in breast cancer CHCs that far exceed previous reports of stem phenotypes within primary tumor tissue [61]. This foundational work sets a path for deeper investigations of the biology underlying tumor-derived hybrid cells and their clinical utility as a circulating biomarker with liquid biopsy potential.

## 4. Materials and Methods

### Human samples and ethics statement

All human peripheral blood and tissue samples were collected and analyzed under approved protocols in accordance with the ethical requirements and regulations of the Oregon Health & Science University (OHSU) institutional review board. Informed consent was obtained from all subjects. Peripheral blood was obtained from cancer patients at OHSU with non-small cell lung cancer, esophageal cancer (ECA), pancreatic ductal adenocarcinoma (PDAC), pancreatic neuroendocrine tumor (PNET), breast cancer, ovarian carcinoma, colon cancer, rectal cancer, adult and pediatric glioma, uveal melanoma, head and neck squamous cell carcinoma (HNSCC), cholangiocarcinoma, ampullary carcinoma, and healthy subjects (n = 5 each site, Table 1, Table S1). Additional specimen from n = 27 patients with untreated breast cancer and n=1 patient with relapsed refractory breast cancer were analyzed (Table 2, Table S2).

### Flow cytometric analyses of CHCs and CTCs in human peripheral blood

Patient peripheral blood was collected in heparinized vacutainer tubes (BD Biosciences, CA) and diluted 1:2 with PBS. Peripheral blood mononuclear cells (PBMCs) were isolated either using RBC lysis or using density centrifugation with Ficoll-Paque PLUS (GE Healthcare, IL). RBC lysis was performed with a 45-minute incubation with Dextran T500 (Pharmacosmos, Denmark, PBS, 3% Dextran 0.1% Sodium Azide), then the top fraction was centrifuged, and the pellet subjected to a 1 minute incubation in 0.2% NaCl followed by addition of the equivalent volume of 1.6% NaCl). Density centrifugation was performed by adding 12ml Ficoll at the bottom of a conical tube containing 20mls of blood and PBS and centrifuging for 20 minutes at 800g with no brake. Isolated PBMCs were counted on a Countess Automated Cell Counter (Thermo Fisher Scientific, MA) and resuspended in FACS Buffer (PBS, 1.0 mM EDTA, 2% FBS) to a concentration of 5×10^7^ cells/mL and 10^7^ cells were then prepared for antibody staining. All staining for flow cytometry was completed on ice, and cells were pelleted with centrifugation at 300 g for 5 minutes after each step. Cells were incubated in PBS containing Live Dead Aqua (Thermo Fisher Scientific, MA) with Fc Receptor Binding Inhibitor (Thermo Fisher Scientific, MA) for 20 minutes. Cells were then incubated in FACS buffer for 30 minutes on ice with CD45-APC or PE (Thermo Fisher Scientific, MA), EPCAM-FITC (Abcam, United Kingdom) or NKI-Beteb-AF647 (Novus Biologicals, CO). For evaluation of stem properties, cells were incubated with EpCAM-BV421 (Abcam, United Kingdom), CD45-FITC (Biolegend, CA), CD31-FITC (BD Biosciences, CA), CD44-PE/Cy7 (BD Biosciences, CA), and CD24-PE (Biolegend, CA), or EpCAM-FITC (Abcam, United Kingdom), CD44-APC (Biolegend, CA), CD24-APC/Cy7 (Biolegend, CA), and CD45-A647 (Biolegend, CA). To prepare cells for intracellular staining, they were incubated with eBioscience Fixation/Permeabilization solution (Thermo Fisher Scientific, MA) for 30 minutes and washed with eBioscience Permeabilization Buffer (Thermo Fisher Scientific, MA). Cells were incubated in 200 μL Permeabilization Buffer with pan-Cytokeratin-PE antibody (Abcam, United Kingdom) for 30 minutes (Table S3). A BD LSRFortessa and Aria Fusion (Becton Dickinson, NJ) FACS machine was used for sample analyses. The gates were established with single color and unstained controls (Figure S1, S3, S4).

### ddPCR detection of KRAS Mutations

After PBMCs were isolated from the peripheral blood of a patient with PDAC, circulating cell populations were collected by FACS. Utilizing the same methodology described for flow cytometery, PBMCs were stained with CD45-APC/Cy7 (Biolegend, CA), EpCAM-FITC (Abcam, United Kingdom), ECAD-PE (Cell signaling, MA), and ITGA3-APC (Biolegend, CA). Using a BD FACSAria™ Fusion (BD Biosciences, CA) cell sorter, 580 CHCs and 7000 normal leukocytes were isolated into FACS buffer. CHCs were defined by CD45 positivity and staining of any tumor marker (EpCAM, ECAD, ITGA3) either in singularly or in combination, while normal leukocytes were defined as CD45+/ECAD^−^/EpCAM^−^/ITGA3^−^. To achieve minimal cell requirements for DNA extraction, 2000 normal leukocytes were spiked into the CHC sample, leaving 5,000 cells in the normal leukocyte sample. All samples were equilibrated to a total volume of 50 μL and then DNA was extracted using Zymo Quick-DNA Microprep Kit (Zymo Research, CA). Procedure was performed using modifications to the manufacturer’s protocol. Briefly, samples were incubated in proteinase K for a minimum of 30 minutes, 15 μLs of the sample were heated to 65 °C and applied to the extraction column, followed by a 10-minute incubation, repeated twice. ddPCR was performed using the ddPCR™ *KRAS* G12/G13 Screening Kit #1863506 (Bio-Rad Laboratories, CA) which detects 7 *KRAS* mutations (*KRAS* p.G12A, p.G12C, p.G12D, p.G12R, p.G12S, p.G12V, and p.G13D), as well as wild type *KRAS*. Droplets were generated with the Auto Droplet Generator (Bio-Rad Laboratories, CA) and measured on the QX200™ Droplet Reader (Bio-Rad Laboratories, CA). Manufacturer recommended parameters for PCR were followed (95 °C for 10 min, followed by 40 cycles of 94 °C for 30 s and 55 °C for 1 min, followed by a final 98 °C heat treatment of 10 minutes for enzyme deactivation). Mutant and wild type *KRAS* thresholds were set to ≥99% of mutant (A549 cell line) and wild type (A375 cell line) controls were positive and 100% of buffer alone specimens were negative (Table S4).

### In situ detection and quantification of CHCs and CTCs from human peripheral blood

PBMCs were isolated from peripheral blood using density centrifugation with Ficoll-Paque PLUS as previously described, and resuspended in FACS Buffer. Cells were then adhered to poly-D-lysine–coated slides through incubation at 37 °C for 15 minutes, permeabilized with Triton-X, and fixed with 4% PFA. Slides were stained with antibodies directed at cancer-specific antigens and CD45 (Table S3) and with DAPI. Slides were imaged using a Zeiss AxioObserver.Z1 light microscope, digitally scanned with a Zeiss AxioScanner.Z1, and analyzed using Zeiss Zen blue software (Carl Zeiss AG, Germany.)

Manual quantification was performed for randomly selected slide regions containing >50,000 nuclei by individuals blinded to the clinical status of the patients or healthy controls. Thresholds for positivity were set off histograms of the unstained portions of the slides. Cells with DAPI nuclear staining were evaluated for CD45, CK, GFAP, CHGA/SYP status. At least 50,000 cells per patient were enumerated. CTCs were defined as tumor-marker (CK, GFAP, CHGA/SYP) positive, CD45 negative. CHCs were defined as cells with both tumor-marker and CD45 staining. Enumerated cells were normalized to 50,000 nuclei.

### Statistical analysis of enumerated CHCs and CTCs

Fisher’s exact test was performed for comparisons of CHC and CTC proportions for each specific disease site and between patients and healthy controls. Additional analyses were performed using Fisher’s one-tailed t-test, using CHC and CTC levels normalized to 500,000 live cells (for flow cytometry) and 50,000 nuclei (for immunohistochemistry) as numbers for each patient or control. Fischer’s exact test was chosen as the primary test because values of zero do not change with normalization. All analyses were performed using SPSS 26 (IBM, New York).

### Mice

All mouse experiments were performed in accordance to the guidelines issued by the Animal Care and Use Committee at Oregon Health & Science University using approved protocols. Mice were housed in a specific pathogen-free environment under strictly controlled light cycle conditions, fed a standard rodent Lab Chow (#5001 PMI Nutrition International, St. Louis, MO), and provided water ad libitum. The following strains were used in the described studies: C57BL/6J (JAX #000664), FVB/N-Tg(MMTV-PyVT)634Mul/J (MMTV:PyMT; JAX #022974)[62]backcrossed onto C57BL/6J for 10 generations, B6.Cg-Tg(CAG-mRFP1)1F1Hadj/J (Cag-RFP; JAX #005884) [63] and Tg(act-EGFP)Y01Osb (Act-GFP; JAX #006567) [64]. Female mice were exclusively used for this study.

### Generation of in vivo-derived hybrid cells

For isolation of *in vivo*-derived hybrids for assessment of hybrid cell tumor-initiating capacity, MMTV:PyMT mice were crossed onto a Cag-RFP background to yield PyMT:RFP mice. Female PyMT:RFP mice were allowed to age until mammary tumors measured 1 cm^3^, approximately 100 days. Tumors were resected, diced, and digested for 30 minutes at 37 °C in DMEM + 2 mg/mL Collagenase A (Roche, Basel, Switzerland) + DNase (Roche) under stirring conditions. Digested tumor was filtered through a 40 μm filter and washed with PBS. Dissociated single cells (1×10^6^) were injected into the mammary fat pad of recipient Act-GFP mice (n=9). Tumors were allowed to grow until they reached a range of 1-2 cm^3^ in volume. Tumors were dissected and processed for FACS-isolation of hybrid cells, or a small specimen of each tumor processed for immunohistochemical analyses, as previously described. Tissue sections were stained with antibodies to E-cadherin to identify neoplastic hybrids within the tumor.

### Analyses of tumor-initiating capacity of hybrids and unfused tumor cells

For FACS-isolation of hybrid cells and unfused tumor cells, tumors were harvested as described above, dissociated to single cells, and subjected to FACS-isolation by direct fluorescence on a Becton Dickinson InFlux sorter. To assess tumor-initiating capacity, 2,500 double-positive RFP^+^/GFP^+^ cells (hybrids), or singly-positive RFP^+^/GFP^−^ cells (unfused tumor cells) were injected into the mammary fat pad of wild type recipient mice (technical replicates, n = 3, each). Tumor growth was monitored and measured when tumors became palpable. Mice were sacrificed when tumor reached 2 cm^3^ in diameter. A second round of tumor cell injections were conducted with 250 hybrid cells or 25,000 and 250,000 unfused tumor cells (n = 3-4).

### Immunohistochemical analyses of tumor tissue

Formalin-fixed paraffin-embedded (FFPE) human tissues were sectioned at 4-5 microns and mounted on charged slides (Tanner Adhesive Slides, Mercedes Medical, TNR WHT45AD). The slides were baked overnight in a Robbin Scientific oven at 55° C and an additional 30 minutes at 65 °C. Tissues were deparaffinized with xylene and rehydrated with graded ethanol baths. Two step antigen retrieval was performed in the Biocare Medical Decloaking Chamber Pro using the following settings: set point 1 (SP1), 125 °C, 30 seconds; SP2: 90 °C, 30 seconds; SP limit: 10 °C. Slides were further incubated in hot pH 9 buffer for 15 minutes. Slides were then washed in two brief changes of diH2O (~2 seconds) and once for 5 minutes in 1x phosphate buffered saline (PBS), pH 7.4 (Fisher, BP39920). Sections were blocked in 10% normal goat serum (NGS, Vector S-1000), 1% bovine serum albumin (BSA, Sigma A7906) in PBS for 30 minutes at 20 °C in a humid chamber, followed by PBS washes.

Primary antibodies (supplemental table 3) were diluted in 5% NGS, 1% BSA in 1x PBS and applied overnight at 4° C in a humidity chamber, covered with plastic coverslips (IHC World, IW-2601). Following overnight incubation, tissues were washed 3 x 10 min in 1x PBS. Coverslips (Corning; 2980-243 and 2980-245) were mounted in Slowfade Gold plus DAPI mounting media (Life Technologies, S36938).

### Fluorescence microscopy

Fluorescently stained slides were scanned on the Zeiss AxioScan.Z1 (Zeiss, Germany) with a Colibri 7 light source (Zeiss). The filter cubes used for image collection were DAPI (Semrock, LED-DAPI-A-000), AF488 (Zeiss 38 HE), AF555 (Zeiss 43 HE), AF647 (Zeiss 50) and Alexa Fluor 750 (AF750, Chroma 49007 ET Cy7). The exposure time was determined individually for each slide and stain to ensure good dynamic range but not saturation. Full tissue scans were taken with the 20x objective (Plan-Apochromat 0.8NA WD=0.55, Zeiss) and stitching was performed in Zen Blue image acquisition software (Zeiss).

### Quenching fluorescence signal

After successful scanning, slides were soaked in 1x PBS for 10 – 30 minutes in a glass Coplin jar, waiting until glass coverslip slid off without agitation. Quenching solution containing 20 mM sodium hydroxide (NaOH) and 3% hydrogen peroxide (H2O2) in 1 x PBS was freshly prepared from stock solutions of 5 M NaOH and 30% H2O2, and each slide placed in 10 ml quenching solution. Slides were quenched under incandescent light, for 30 minutes for FFPE tissue slides and 20 minutes for PBMCs adhered to glass slides. Slides were then removed from chamber with forceps and washed three times for two minutes in 1 x PBS. The next round of primary antibodies was applied, diluted in blocking buffer as previously described, and imaging and quenching were repeated over ten rounds for FFPE tissue slides, and two rounds for PBMCs adhered to glass slides.

### Digital quantification and analysis of FFPE tissue cyclic immunofluorescence

Each image acquired during the cyCIF assay was registered based on DAPI features acquired from each round of staining [65]. In-house software [66] was used to generate nuclear, cell and membrane segmentation masks by classifying pixels on the basis of a combination of marker expression to identify cells and membranes respectively. Extracted single-cell features included centroids and mean intensity of each marker from its biologically-relevant segmentation mask, e.g. Ecad_Ring, Ki67_Nuclei. The last round DAPI image was used to filter out cells lost during each round of cyCIF staining. For downstream analysis, first intensity normalization is performed for each biopsy sample based on RESTORE (robust intensity normalization method) [67] to minimize intensity variation across samples. Then, a heatmap was constructed using unsupervised clustering [k-means clustering with the number of clusters (n=20)].

### Quantification and analysis of PBMC cyclic immunofluorescence

Digitally scanned slides as described above were processed using Zeiss blue software. Regions of interest were created and conserved between round of staining and quenching. Thresholds for positivity were set off histograms of the unstained portions of the slides. Region of interest were registered using the image correlation feature, registered cell overlays were used for phenotypic profiling at the single cell level. Cells with DAPI nuclear staining were evaluated for CD45, CK, ECAD, ER, AR, Ki67 and CD44 status. At least 50,000 cells were enumerated. CTCs were defined as cells with positive tumor-marker staining with no CD45 staining compared to other PBMCs. CHCs were defined as cells with both tumor-marker staining and strong CD45 staining.

## 5. Conclusions

In conclusion, we demonstrate that immune-neoplastic cell hybrids represent a heterogeneous population of cells that disseminate into circulation as CHCs and is a generalizable phenomenon in human solid malignancies. Neoplastic hybrids harbor genetic hallmarks of parental tumor cells, display stem markers, and recapitulate tumorigenesis to a greater degree than unfused cancer cells. Finally, we reveal that CHC phenotypes reflect the heterogeneity of functional protein expression from cancer tissue. This novel population is deserving of further study in an effort to understand the extent to which it can be leveraged as a liquid biomarker and in treatments to forestall metastatic progression.

## Supplementary Materials

The following are available online at www.mdpi.com/xxx/s1 Figure S1: Disseminated tumor cell populations by flow cytometry, Figure S2: Disseminated tumor cell populations by immunofluorescence microscopy, Figure S3: Murine tumor cell populations by flow cytometry, Figure S4: Phenotype characterization of disseminated tumor cells by flow cytometry. Table S1: Clinicopathologic characteristics of individual patients with assorted malignancies, Table S2: Clinicopathologic characteristics of individual breast cancer patients evaluated for stem properties of circulating tumor-derived cells, Table S3: Antibodies and reagents, Table S4: Digital droplet PCR results and controls.

## Author Contributions

Conceptualization, MS Dietz, TL Sutton, BS Walker, CE Gast, L Zarour, and MH Wong; methodology, MS Dietz, TL Sutton, BS Walker, CE Gast, J Eng, K Chin, E Burlingame, and YH Chang; formal analysis, MS Dietz, TL Sutton, BS Walker, CE Gast, E Burlingame, YH Chang, J Eng, K Chin, and MH Wong; investigation, MS Dietz, TL Sutton, BS Walker, CE Gast, L Zarour, JR Swain, J Eng, M Parappilly, K Limbach, A Sattler, E Burlingame, A Sapre, JL Montoya Mira, A Gower, and Y Chin; resources, JM Mira, A Sapre, Y Chiu, DR Clayburgh, SJ Pommier, JP Cetnar, JM Fischer, K Tolba, JJ Jaboin, SJ Han, KJ Nazemi, RF Pommier, GK Billingsley, BC Sheppard, AH Skalet, VL Tsikitisi, SC Mayo, CD Lopez, JW Gray, G Mills and Z Mitri; data curation, SK Sengupta; writing—original draft preparation, MS Dietz, TL Sutton, BS Walker, and MH Wong; writing—review and editing, JP Dolan, AH Skalet, MS Dietz, TL Sutton, BS Walker, and MH Wong; visualization, BS Walker, SK Sengupta, MH Wong, and MS Dietz,; supervision, MS Dietz, TL Sutton, BS Walker, CE Gast, K Chin, YH Chang, and MH Wong; funding acquisition, YH Chang, and MH Wong. All authors have read and agreed to the published version of the manuscript.

## Acknowledgments

We acknowledge members of the OHSU Flow Cytometry Shared Resource, and Advanced Light Microscopy Core at The Jungers Center for technical assistance. We also acknowledge support from the OHSU Cancer Early Detection and Research (CEDAR) department for patient identification.

