## Supplemental data for "Relevance of Circulating Hybrid Cells as a Non-Invasive Biomarker for Myriad Solid Tumors"

**Supplementary Tables and Figures**

**Supplementary Figure 1:** Gating Scheme for Identification of CHCs and CTCs by Flow Cytometry

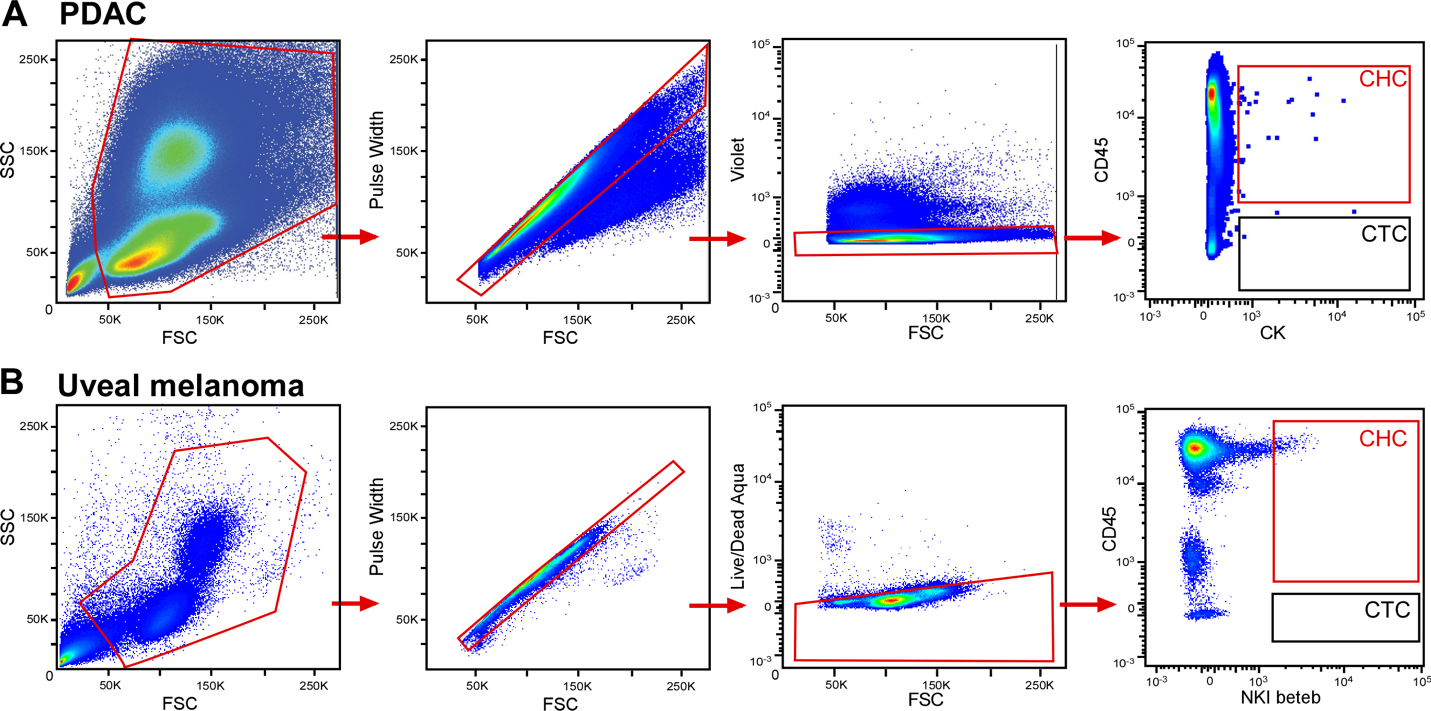

**Supplementary Figure 1:** Isolated human peripheral blood mononuclear cells from patients with **A)** epithelial cancers and **B)** uveal melanoma were stained and subjected to FACS or analysis by flow cytometry. Gating scheme established based upon single color controls and/or fluorescence minus one control. Representative data from Fig. 1B.

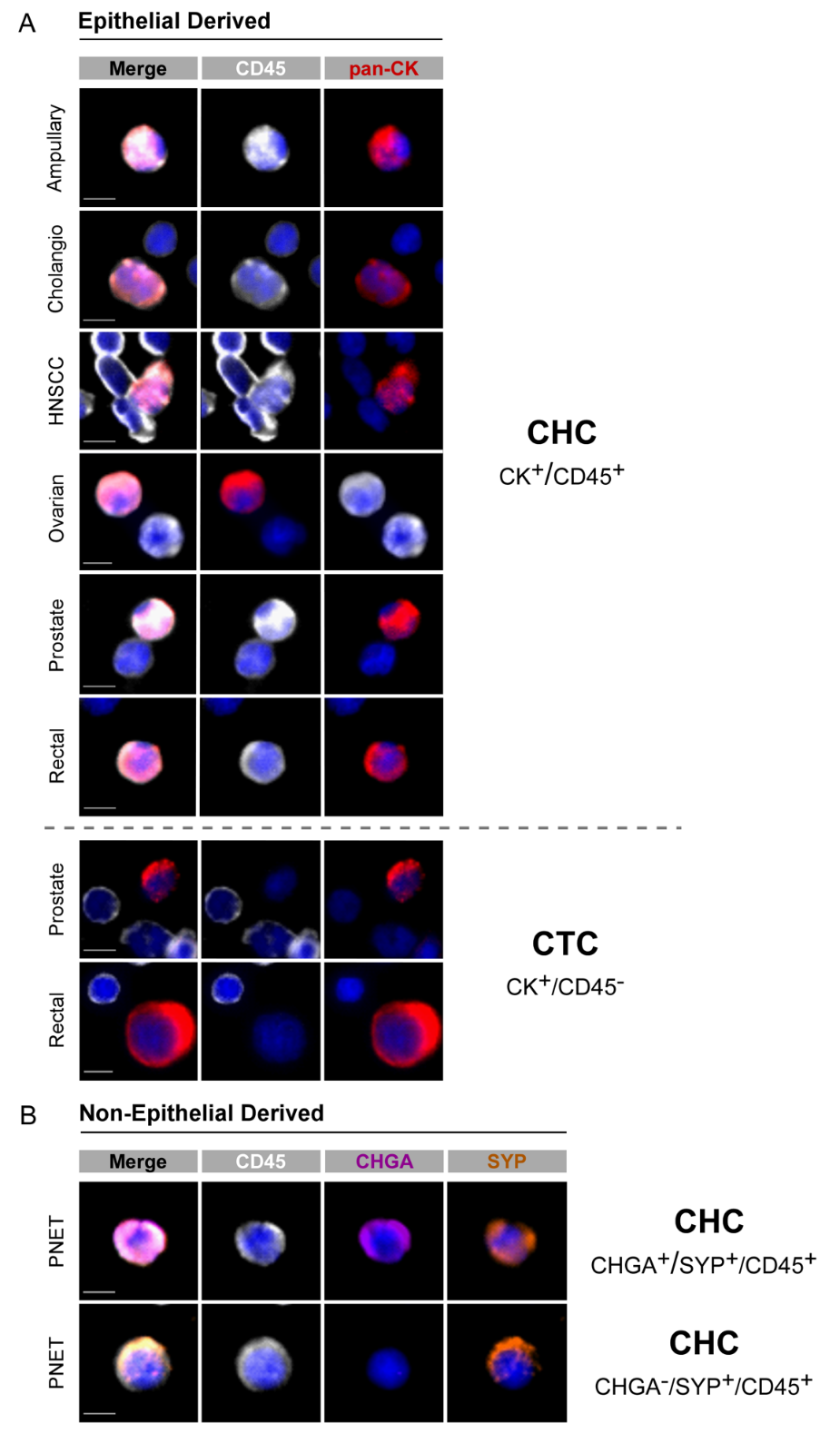
**Supplementary Figure 2:** Tumor-derived Cells in Circulation Identified in Human Patients by Immunofluorescence Microscopy

**Supplementary Figure 2:** Disseminated tumor-derived cells identified in peripheral blood samples from human patients with **A)** epithelial and **B)** non-epithelial cancers using immunofluorescence microscopy. CHCs from epithelial cancers are defined by dual expression of cytokeratin (CK) and CD45, while CTCs are CK^+^/CD45^-^. Nuclei are counterstained with DAPI (blue). Pancreatic neuroendocrine tumor (PNET)-derived CHCs express CD45 along with chromogranin A (CHGA) and synaptophysin (SYN), either individually or in combination. Scale bar: 5 µm

**Supplementary Figure 3:** Gating Scheme for Isolating Murine-Derived Fused and Unfused Tumor Cells

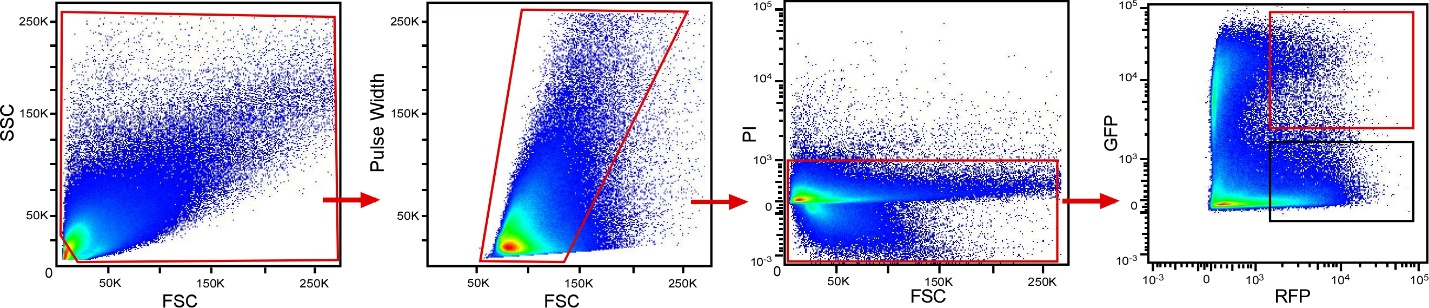

**Supplementary Figure 3:** Dissociated tumor cells were stained and subjected to FACS. Gating scheme established based upon single color controls and/or FMO controls. RFP-expressing unfused tumor cells and fusion hybrids co-expressing RFP and GFP were isolated for injection into mice. Representative data from Fig. 2B

**Supplementary Figure 4:** Gating Scheme for Identification of CHC and CTC Subpopulations by Flow Cytometry

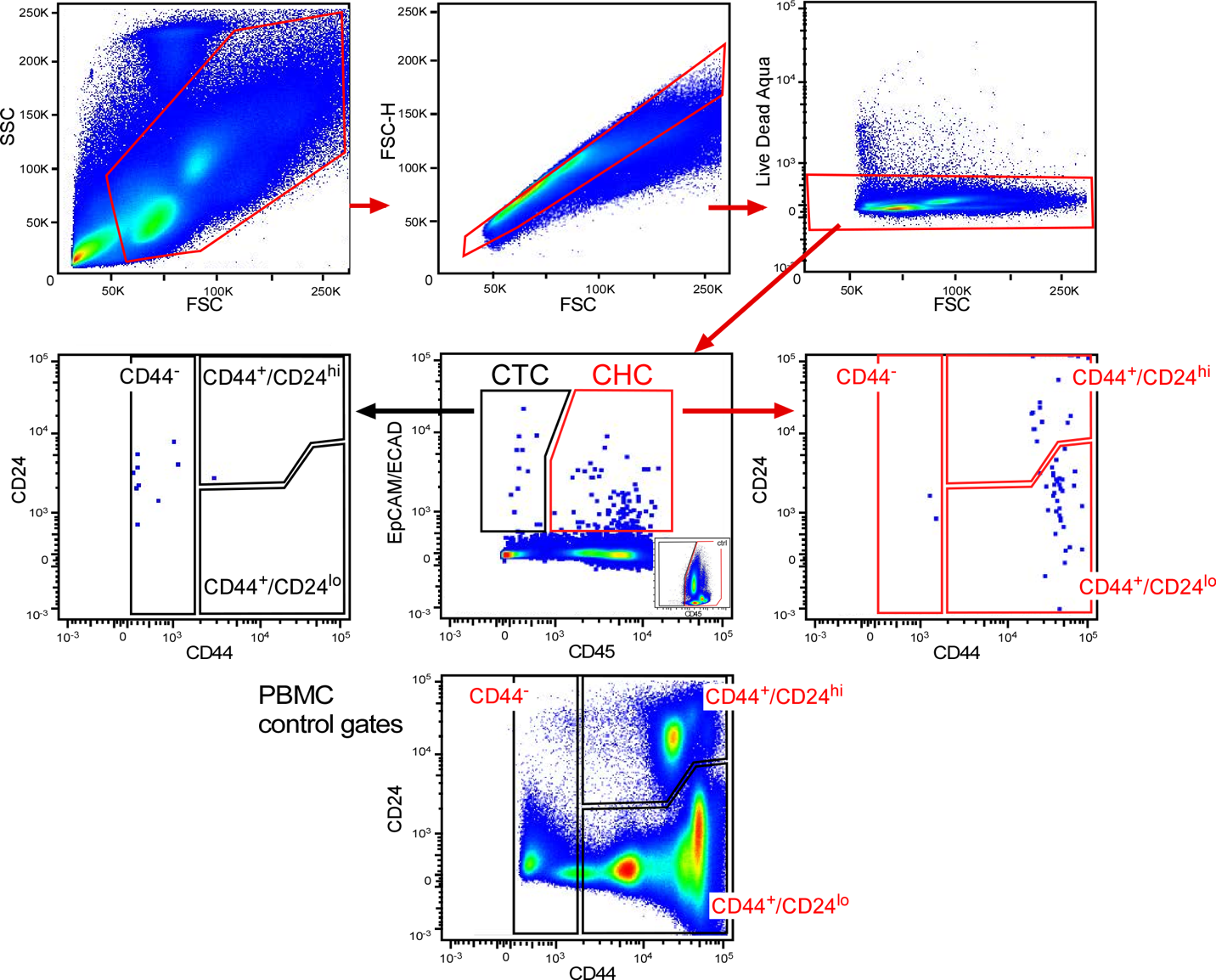

**Supplementary Figure 4:** Isolated human peripheral blood mononuclear cells from patients with breast cancer were stained and analyzed by flow cytometry. Gating scheme established based upon unstained controls. Representative data from Fig. 3.

Supplementary Table 1: Clinicopathologic Characteristics of Individual Patients with Assorted Malignancies

| **Patient ID** | **Age** | **Sex** | **Site-Specific Characteristic** | **Disease Burden** | **Prior Treatment*** | **Vital Status** |
| --- | --- | --- | --- | --- | --- | --- |
| **Ampullary Adenocarcinoma** |  |  | *Subtype* |  |  |  |
| A1 | 77 | F | Intestinal | Local | None | NED |
| A2 | 74 | M | Intestinal | Regional | None | DOD |
| A3 | 62 | F | Pancreaticobiliary | Regional | None | DOD |
| A4 | 73 | F | Pancreaticobiliary | Regional | None | NED |
| A5 | 56 | F | Panctreaticobiliary/Intestinal Mixed | Local | None | NED |
| **Breast Adenocarcinoma** |  |  | *Hormone Receptor and HER2 Status* |  |  |  |
| B1 | 73 | F | HR+/HER2- | Regional | None | NED |
| B2 | 73 | F | HR+/HER2- | Local | None | NED |
| B3 | 52 | F | HR+/HER2- | Regional | None | AWD |
| B4 | 51 | F | HR-/HER2+ | Metastatic | None | AWD |
| B5 | 70 | F | HR+/HER2- | Metastatic | Pembrolizumab | AWD |
| **Lung Cancer** |  |  | *Histology* |  |  |  |
| L1 | 74 | M | Squamous Cell Carcinoma | Metastatic | Nivolumab | DOD |
| L2 | 41 | F | Squamous Cell Carcinoma | Metastatic | Carboplatin, Paclitaxel | DOD |
| L3 | 70 | M | Adenocarcinoma | Metastatic | None | DOD |
| L4 | 60 | M | Squamous Cell Carcinoma | Metastatic | None | DOD |
| L5 | 53 | M | Adenocarcinoma | Metastatic | Nivolumab | NED |
| **Colon Adenocarcinoma** |  |  | *Mutational Status* |  |  |  |
| C1 | 59 | M | Not tested | Metastatic | None | DOD |
| C2 | 33 | F | MSH6 mutant | Metastatic | FOLFOX, bevacizumab | NED |
| C3 | 65 | M | Wild type** | Metastatic | CapeOx, bevacizumab | DOD |
| C4 | 50 | F | KRAS mutant | Metastatic | FOLFOX | DOD |
| C5 | 57 | F | Wild type** | Metastatic | FOLFOX, panitumumab | AWD |
| **Rectal Adenocarcinoma** |  |  | *Mutational Status* |  |  |  |
| R1 | 67 | F | Wild type ** | Metastatic | FOLFOX | AWD |
| R2 | 50 | F | Wild type ** | Metastatic | FOLFOX | DOD |
| R3 | 63 | M | Wild type ** | Metastatic | FOLFOX | NED |
| R4 | 47 | F | KRAS mutant | Metastatic | FOLFOX | DOD |
| R5 | 49 | M | KRAS mutant | Metastatic | FOLFOX | NED |
| **Esophageal Cancer** |  |  | *Histology* |  |  |  |
| E1 | 52 | M | Adenocarcinoma | Regional | Carboplatin, Paclitaxel, Radiation | DOD |
| E2 | 70 | M | Adenocarcinoma | Regional | Carboplatin, Paclitaxel, Radiation | DOD |
| E3 | 75 | M | Adenocarcinoma | Local | Carboplatin, Paclitaxel, Radiation | NED |
| E4 | 53 | M | Squamous Cell Carcinoma | Local | Carboplatin, Paclitaxel, Radiation | DOD |
| E5 | 66 | M | Adenocarcinoma | Local | Carboplatin, Paclitaxel, Radiation | DOD |
| **Ovarian Carcinoma** |  |  | *Histology* |  |  |  |
| OVA1 | 75 | F | High-Grade Serous Cacinoma | Regional | None | LTF |
| OVA2 | 61 | F | High-Grade Serous Cacinoma | Regional | Carboplatin, Paclitaxel | AWD |
| OVA3 | 68 | F | High-Grade Serous Cacinoma | Metastatic | Carboplatin, Paclitaxel | NED |
| OVA4 | 76 | F | High-Grade Serous Cacinoma | Regional | Carboplatin, Paclitaxel | NED |
| OVA5 | 60 | F | High-Grade Serous Cacinoma | Regional | Carboplatin, Paclitaxel | NED |
| **Pancreatic Adenocarcinoma** |  |  | *Histology* |  |  |  |
| PDAC1 | 75 | M | Adenocarcinoma | Metastatic | FOLFIRINOX | AWD |
| PDAC2 | 72 | F | Adenocarcinoma | Regional | FOLFIRINOX | NED |
| PDAC3 | 76 | M | Adenocarcinoma | Regional | FOLFIRINOX | NED |
| PDAC4 | 66 | F | Adenocarcinoma | Metastatic | None | AWD |
| PDAC5 | 56 | F | Adenocarcinoma | Local | Olaparib | NED |
| **Pancreatic NET** |  |  | *WHO Grade/Functional Status* |  |  |  |
| PNET1 | 55 | M | G1/Nonfunctional | Local | None | NED |
| PNET2 | 65 | F | G1/Insulinoma | Local | None | NED |
| PNET3 | 88 | F | G2/Nonfunctional | Regional | None | NED |
| PNET4 | 64 | F | G1/Nonfunctional | Local | None | NED |
| PNET5 | 59 | M | G3/Nonfunctional | Local | None | NED |
| **Uveal Melanoma** |  |  | *GEP Class* |  |  |  |
| U1 | 67 | M | 2, PRAME unknown | Metastatic | None | AWD |
| U2 | 56 | M | 2, PRAME+ | Local | None | NED |
| U3 | 71 | M | 2, PRAME+ | Local | None | DOD |
| U4 | 59 | F | 2, PRAME+ | Local | None | NED |
| U5 | 67 | M | 2, PRAME+ | Local | None | NED |
| **Head & Neck SCC** |  |  | *Primary Location* |  |  |  |
| H1 | 60 | M | Buccal | Regional | None | LTF |
| H2 | 70 | M | Floor of Mouth | Regional | None | LTF |
| H3 | 57 | F | Floor of Mouth | Regional | None | DOD |
| H4 | 69 | F | Oral Tongue | Regional | None | NED |
| H5 | 63 | F | Floor or Mouth | Regional | None | NED |
| **Adult Glioblastoma** |  |  | *Mutational Status* |  |  |  |
| AG1 | 31 | M | IDH WT, MGMTP nonmethylated, H3 K27M mutant | Local | Radiation, temozolamide | AWD |
| AG2 | 74 | M | IDH WT, EGFR3 mutant, MGMTP nonmethylated | Local | None | DOD |
| AG3 | 67 | F | IDH status unknown, MGMTP hypermethylated | Local | Carmustine wafer implantation | AWD |
| AG4 | 68 | F | IDH WT, MGMTP nonmethylated | Local | Temozolamide | AWD |
| AG5 | 47 | M | EGFR3 mutant, IDH WT, MGMTP nonmethylated | Local | None | AWD |
| **Pediatric Glioma** |  |  | *Subclass* |  |  |  |
| PG1 | 6 | F | Diffuse Midline Glioma | Local | Radiation, Temozolamide | DOD |
| PG2 | 4 | F | Diffuse Midline Glioma | Local | Panobinostat | DOD |
| PG3 | 6 | F | Diffuse Midline Glioma | Local | Panobinostat | DOD |
| PG4 | 5 | M | Grade III Anaplastic Astrocytoma | Local | Radiation | AWD |
| PG5 | 15 | M | Glioblastoma- EGFR Amplified | Local | None | AWD |
| **Prostate Adenocarcinoma** | |  | *Gleason Score* |  |  |  |
| P1 | 60 | M | 3 + 4 = 7 | Local | None | AWD |
| P2 | 69 | M | 3 + 3 = 6 | Local | None | NED |
| P3 | 72 | M | 3 + 3 = 6 | Local | None | NED |
| P4 | 57 | M | 3 + 4 = 7 | Local | None | NED |
| P5 | 61 | M | 3 + 4 = 7 | Local | None | AWD |
| **Cholangiocarcinoma** |  |  | *Anatomic Location* |  |  |  |
| CC1 | 42 | F | Distal Common Bile Duct | Metastatic | None | AWD |
| CC2 | 65 | F | Distal Common Bile Duct | Local | None | NED |
| CC3 | 81 | M | Distal Common Bile Duct | Node Positive | None | AWD |
| CC4 | 78 | F | Proximal Common Bile Duct | Metastatic | None | DOD |
| CC5 | 60 | M | Intrahepatic Bile Duct | Local | None | NED |
| **Normal Controls** |  |  | *History of Cancer or Chronic Inflammatory Diseases* | *Paradigm* |  |  |
| N1 | 23 | F | No | Flow | - | - |
| N2 | 28 | M | No | Flow | - | - |
| N3 | 25 | M | No | Flow | - | - |
| N4 | 24 | M | No | Flow | - | - |
| N5 | 21 | M | No | Flow | - | - |
| N6 | 31 | M | No | Flow + IF | - | - |
| N7 | 24 | F | No | Flow + IF | - | - |
| N8 | 26 | F | No | Flow + IF | - | - |
| N9 | 25 | F | No | Flow + IF | - | - |
| N10 | 23 | M | No | Flow + IF | - | - |
| N11 | 81 | M | No | IF | - | - |
| N12 | 65 | M | No | IF | - | - |
| N13 | 73 | F | No | IF | - | - |
| N14 | 67 | F | No | IF | - | - |
| N15 | 64 | F | No | IF | - | - |
| * Any chemotherapy or immunotherapy within 8 weeks, any curative-intent surgery, or radiotherapy within 3 months of sample collection ** Wild type with reference to KRAS, BRAF, NRAS, and mismatch repair genes (MLH1, MSH2, MSH6 and PMS2) Abbreviations: HR=Hormone Receptor; HER2=Human Epidermal Growth Factor Receptor 2; DOD=Dead of Disease; FOLFOX=5-Fluorouracil, folinic acid, oxaliplatin; NED=No evidence of disease; FOLFIRI=5-FU, folinic acid, irinotecan; LTF=Lost to Followup; AWD= Alive with Disease; M=Male; F=Female; CapeOx=Capecitabine, Oxaliplatin; MGMTP=O6-methylguanine–DNA methyltransferase promoter; IDH=Isocitrate dehydrogenase; EGFR3=Epidermal Growth Factor Receptor 3; PRAME=Preferentially Expressed Antigen in Melanoma; GEP= Gene Expression Profiling; SCC=Squamous Cell Carcinoma; WHO=World Health Organization; NET= Neuroendocrine tumor | | | | | | |

Supplementary Table 2: Clinicopathologic Characteristics of Individual Breast Cancer Patients Evaluated for Stem Properties of Circulating Tumor-Derived Cells

| **Patient** | | **Age** | **Sex** | **HR Status** | **HER2 Status** | **TNBC** | **Grade** | **Staging*** | **Prior Treatment**** | **Vital Status** |
| --- | --- | --- | --- | --- | --- | --- | --- | --- | --- | --- |
|  | Breast 1 | 66 | F | + | + | - | III | pT2N0M0 | None | NED |
|  | Breast 2 | 22 | F | + | - | - | II | pT1N0M0 | None | NED |
|  | Breast 3 | 28 | F | + | - | - | II | pT2N1M0 | None | AWD |
|  | Breast 4 | 79 | F | + | - | - | III | pT2N2M0 | None | NED |
|  | Breast 5 | 59 | F | + | + | - | III | pT3N1M1 | None | DOD |
|  | Breast 6 | 44 | F | - | + | - | III | cT3N3M0 | None | NED |
|  | Breast 7 | 54 | F | + | - | - | II | pT1N0M0 | None | NED |
|  | Breast 8 | 76 | F | + | - | - | I | pT1cN0M0 | None | NED |
|  | Breast 9 | 63 | F | + | - | - | II | pT2N0M0 | None | NED |
|  | Breast 10 | 38 | F | + | + | - | II | cT2N0M0 | None | NED |
|  | Breast 11 | 72 | F | + | - | - | II | pT2N1M0 | None | NED |
|  | Breast 12 | 58 | F | - | - | + | III | cT2N2M0 | None | NED |
|  | Breast 13 | 77 | F | + | + | - | III | pT1N1M0 | None | NED |
|  | Breast 14 | 67 | F | + | + | - | III | cT2N1M0 | None | NED |
|  | Breast 15 | 80 | F | + | - | - | II | pT4bN3M1 | None | DOD |
|  | Breast 16 | 44 | F | + | - | - | II | pT2N0M0 | None | NED |
|  | Breast 17 | 28 | F | + | - | - | II | cT1N0M0 | None | NED |
|  | Breast 18 | 60 | F | + | - | - | III | pT2N1M0 | None | NED |
|  | Breast 19 | 75 | F | + | - | - | II | pT1NxM0 | None | NED |
|  | Breast 20 | 55 | F | + | - | - | II | pT2N0M0 | None | NED |
|  | Breast 21 | 26 | F | - | - | + | III | cT3N1M0 | None | AWD |
|  | Breast 22 | 76 | F | + | + | - | III | cT4N3M1 | None | AWD |
|  | Breast 23 | 55 | F | + | + | - | III | pT1N0M0 | None | NED |
|  | Breast 24 | 52 | F | - | - | + | III | cT2N0M0 | None | NED |
|  | Breast 25 | 52 | F | + | - | - | I | cT3N1M0 | None | NED |
|  | Breast 26 | 52 | F | + | - | - | II | cT1N1M0 | None | AWD |
|  | Breast 27 | 50 | F | - | + | - | III | pT2N1M1 | None | AWD |
| * Staging reported as clinical stage for treatment-naive patients sampled prior to NAC ** Any chemotherapy or immunotherapy within 6 weeks, any curative-intent surgery, or radiotherapy within 3 months of sample collection Abbreviations: HR=Hormone Receptor; HER2= Human Epidermal Growth Factor Receptor 2; TNBC= Triple Negative Breast Cancer, DOD= Dead of Disease; DWOD =Dead without Disease; NED= No evidence of disease; AWD= Alive with Disease; F= Female; NAC= Neoadjuvant Chemotherapy; TCHP=docetaxel, carboplatin, trastuzumab, pertuzumab; ACT= doxorubicin, cyclophosphamide, paclitaxel; ddACT= Dose dense doxorubicin, cyclophosphamide, paclitaxel; ACTHP: doxorubicin, cyclophosphamide, paclitaxel, trastuzumab, pertuzumab; TCH=docetaxel, carboplatin, trastuzumab | | | | | | | | | | |

Supplementary Table 3: Antibodies and Reagents

| **Target** | **Cyclic IF Round** | **Clone** | **Species** | **Manufacturer** | **Fluorophore** | **Dilution Factor*** |
| --- | --- | --- | --- | --- | --- | --- |
| **Immunocytochemistry** | | | | | | |
| CD45 | 1 | HI30 | Ms | BioLegend | FITC | 100 |
| CD44 | 1 | EPR1013Y | Rb | Abcam | A750 | 200 |
| ER | 1 | EPR4097 | Rb | Abcam | A647 | 50 |
| pan-CK | 1 | AE1/3 | Ms | Invitrogen | eFlour570 | 100 |
| pan-CK | - | AE1/3 | Ms | Novus Biologicals | A750 | 500 |
| ECAD | 2 | 24 E 10 | Ms | CST | A750 | 100 |
| AR | 2 | Polyclonal | Rb | Millipore | A555 | 200 |
| Ki67 | 2 | D3B5 | Rb | CST | A647 | 400 |
| GFAP | - | A5 | Rb | Invitrogen | eFlour570 | 200 |
| Synaptophysin | - | YE269 | Rb | Abcam | A555 | 100 |
| Chromogranin A | - | SPM339 | Ms | Novus Biologicals | A647 | 25 |
| **Cyclic Immunofluorescence on Tumor** | | | | | | |
| PCNA | 1 | PC10 | Ms | CST | A488 | 25 |
| CD8 | 1 | C8/468 | Ms | Abcam | A555 | 50 |
| PD1 | 1 | EPR4877(2) | Rb | Abcam | A647 | 50 |
| CK19 | 1 | A53-B/A2 | Ms | Biolegend | A750 | 100 |
| CK5 | 2 | EP1601Y | Rb | Abcam | A488 | 100 |
| HER2 | 2 | 3B5 | Ms | Thermofisher | A555 | 50 |
| ER | 2 | EPR4097 | Rb | Abcam | A647 | 100 |
| CD45 | 2 | EP322Y | Rb | Abcam | A750 | 50 |
| aSMA | 3 | 1A4 | Ms | Santa Cruz | A488 | 100 |
| CD68 | 3 | KP1 | Ms | Biolegend | A555 | 50 |
| CD4 | 3 | EPR6855 | Rb | Abcam | A647 | 100 |
| Ecad | 3 | EP700Y | Rb | Abcam | A750 | 100 |
| Vimentin | 4 | D21H3 | Rb | CST | A488 | 200 |
| AR | 4 | polyclonal | Rb | Sigma | A555 | 100 |
| CD31 | 4 | EPR3094 | Rb | Abcam | A647 | 100 |
| CD44 | 4 | EPR1013Y | Rb | Abcam | A750 | 200 |
| CK7 | 5 | EPR1619Y | Rb | Abcam | A488 | 200 |
| CK14 | 5 | LL002 | Ms | BioRad | A555 | 200 |
| Ki67 | 5 | D3B5 | Rb | CST | A647 | 400 |
| PgR | 5 | YR85 | Rb | Abcam | A750 | 50 |
| pHH3 | 6 | D2C8 | Rb | CST | A488 | 50 |
| pS6 | 6 | D57 | Rb | CST | A555 | 200 |
| FoxP3 | 6 | 206D | Ms | Biolegend | A647 | 50 |
| CD20 | 6 | EP459Y | Rb | Abcam | A750 | 25 |
| CK17 | 7 | EP1623 | Rb | Abcam | A488 | 100 |
| EGFR | 7 | D38B1 | Rb | CST | A555 | 50 |
| pERK | 7 | Erk1/2 | Rb | CST | A647 | 50 |
| GRNZB | 7 | EPR20129-217 | Rb | Abcam | A750 | 100 |
| H3K27 | 8 | C36B11 | Rb | CST | A488 | 50 |
| PDPN | 8 | D2-40 | Ms | Biolegend | A555 | 25 |
| LaminB2 | 8 | EPR9701(B) | Rb | Abcam | A647 | 200 |
| ColI | 8 | EPR7785 | Rb | Abcam | A750 | 50 |
| CK8 | 9 | EP1628Y | Rb | Abcam | A488 | 200 |
| cPARP | 9 | D64E10 | Rb | CST | A555 | 25 |
| pRB | 9 | EPR17732 | Rb | Abcam | A647 | 50 |
| LaminAC | 9 | 4C11 | Ms | Sigma | A750 | 500 |
| LaminB1 | 10 | EPR8985(B) | Rb | Abcam | A488 | 50 |
| H3K4 | 10 | C42D8 | Rb | CST | A555 | 50 |
| ColIV | 10 | 1042 | Ms | eBioscience | A647 | 50 |
| CD3 | 10 | EP449E | Rb | Abcam | A750 | 50 |
| **Flow Cytometry** | | | | | | |
| CD24 |  | ML5 | Ms | Biolegend | PE, APC/Cy7 | 50, 20 |
| CD31 |  | WM59 | Ms | BD | FITC | 50 |
| CD44 |  | G44-26 | Ms | BD | PE/Cy7 | 100 |
| CD44 |  | IM7 | Rat | Biolegend | APC | 100 |
| CD45 |  | HI30 | Ms | Invitrogen | APC | 25 |
| CD45 |  | HI30 | Ms | BD | FITC | 50 |
| CD45 |  | HI30 | Ms | Biolegend | PE | 100 |
| CD45 |  | 2D1 | Ms | Biolegend | APC/Cy7 | 100 |
| pan-CK |  | C11 | Ms | Abcam | PE | 250 |
| EpCAM |  | 9C4 | Ms | Abcam | FITC | 100 |
| EpCAM |  | 9C4 | Ms | Biolegend | BV421 | 100 |
| ECAD |  | 24E10 | Rb | CST | A488, PE | 50 |
| ITGA3 |  | ASC-1 | Ms | Biolegend | APC | 150 |
| Live Dead Aqua |  | - | - | Life Technologies | - | 500 |
| Fc Binding Inhibitor |  | - | - | eBioscience | - | 200 |
| **Uveal Melanoma** | | | | | | |
| PMEL17/SILV |  | NKI/Beteb | Ms | Novus Biologicals | A647 | 500 |
| CD45 |  | HI30 | Ms | Biolegend | FITC | 100 |
| Live Dead Aqua |  | - | - | Life Technologies | - | 500 |
| Fc Binding Inhibitor |  | - | - | eBioscience | - | 200 |
| **Murine Fluorescence-Activated Cell Sorting** | | | | | | |
| ECAD |  | DECMA-1 | Rat | Sigma | Unconjugated | - |
| CD45 |  | 30-F11 | Rat | Biolegend | PE/Cy7 | 8000 |
| Live Dead Aqua |  | - | - | Life Technologies | - | 500 |
| *From stock concentration. For immunohistochemistry and immunocytochemistry, dilutions are given in μg/mL applied to slides. For flow cyometry, dilutions from stock are given with samples incubated at 5x107 cells/ml.  Abbreviations: IF=Immunofluorescence; CD=Cluster of Differentiation; CST=Cell Signaling Technologies; GFAP=Glial Fibrillary Acidic Protein; CK=Cytokeratin; ECAD=E-cadherin; AR=Androgen Receptor; ER=Estrogen Receptor; PCNA=Proliferating Cell Nuclear Antigen; PD1=Programmed Cell Death Protein 1; HER2=Human Epidermal Growth Factor Receptor 2; aSMA=Smooth Muscle/F-actin; PgR=Progesterone Receptor; PHH3=Phosphohistone H3; PS6=Ribosomal Protein S6; FOXP3=Forkhead Box P3; EGFR=Epidermal Growth Factor Receptor; pERK=Phospho-p44/42 MAPK; GRNZB=Granzyme B; H3K27=Tri-Methyl-Histone H3 (Lys27); PDPN=Podoplanin; Col=Collagen; cPARP=Poly(ADP-ribose) Polymerase Cleavage; pRB=Retinoblastoma Protein; H3K4=Tri-Methyl-Histone H3 (Lys4); EpCAM=Epithelial Cell Adhesion Molecule; PMEL17/SILV=Melanocyte Protein Mel-17; Ms=Mouse; Rb=Rabbit; PE=Phycoryethrin; FITC=Fluorescein Isothiocyanate; APC=Allophycocyanin; BV421=Blue Violet-421 | | | | | | |

Supplementary Table 4: Digital Droplet PCR Results and Controls

| **Sample** | **MUT KRAS Droplets** | **WT KRAS Droplets** | **Total Accepted Droplets** | **Volume (ml)** |
| --- | --- | --- | --- | --- |
| CHC* | 7 | 708 | 10586 | 8.8 |
| CD45^+^ | 0 | 393 | 10080 | 8.8 |
| A549  (MUT control) | 6323 | 8 | 24523 | 17.6 |
| A375  (WT control) | 1 | 3416 | 25994 | 17.6 |
| *Sample contained 22% CHCs and 78% normal CD45+ leukocytes  MUT: Mutant; WT: Wild type | | | | |
